# Spinning disk confocal microscopy with a 25 Megapixel Camera

**DOI:** 10.1101/2025.07.30.667773

**Authors:** Guy M. Hagen, Brian Lewis, Summer Levis, Joseph R. Hamilton, Tristan C. Paul

## Abstract

Spinning disk confocal microscopy enables fast optical sectioning with low phototoxicity but is often inaccessible due to high hardware costs. We present a low-cost solution using a 25 megapixel machine vision CMOS camera (Sony IMX540, FLIR Blackfly S) and a custom-built spinning disk. The system uses a back-illuminated sensor with high quantum efficiency (69% at 525 nm) and low read noise (2.31 electrons). High-resolution images of Thy1-GFP mouse brain slices and H&E-stained rat testis verified performance across 3D tissue volumes. The custom disk, made with 18 µm pinholes (180 µm pitch) on a chrome photomask and mounted to an optical chopper motor, enables stable, near-telecentric imaging. Micro-Manager software integration allows synchronized control of all hardware, which demonstrates that affordable CMOS sensors can potentially replace sCMOS in spinning disk microscopy, offering an open-source, scalable solution for advanced imaging.

## 1. Introduction

Fluorescence microscopy is one of the most important laboratory methods in use today in many areas of biological and biomedical research. Research-grade fluorescence microscopes and their related light sources and detectors are prohibitively expensive in some situations, so finding lower cost alternatives is an important goal. Innovative approaches include open-source hardware designs [1–6], and open-source software for microscope control and data analysis [7–10].

Spinning disk confocal microscopy [11,12] is a camera-based method in optical sectioning fluorescence microscopy which is often used in live cell imaging studies [13], but can be very expensive when considering the commercial systems. The scientific CMOS cameras typically used have high quantum efficiency and very low noise together with fast readout. These cameras have a high performance for spinning disk confocal microscopy but are quite expensive. The spinning disk unit itself is also quite costly when considering the newest designs.

For this work, we followed a recent design for a low cost, home-built spinning disk confocal microscope [14] with a few modifications as described below. For detection we used a CMOS camera intended for machine vision purposes. Previous machine vision cameras in this category were less suitable for low light imaging conditions such as those present in fluorescence microscopy, but newer CMOS cameras have made large improvements in performance characteristics necessary for this application. Newer CMOS image sensors have improved quantum efficiency and reduced noise, which together offer large improvements in sensitivity and usefulness for this application.

## 2. Materials and methods

### 2.1 Samples

We imaged an optically cleared coronal mouse brain slice in which a subset of neurons express green fluorescent protein (GFP). The slice was approximately 150 μm thick and was obtained from SunJin Lab (Hsinchu City, Taiwan). This supplier used a Thy1-GFP mouse strain [15], which the supplier prepared as follows:

1. cardiac perfusion with cold, freshly prepared 4% paraformaldehyde (PFA),
2. fixation of the dissected brain with a 4% PFA solution on an orbital shaker overnight at 4 °C followed by washing three times with phosphate-buffered saline (PBS) at room temperature,
3. sectioning the brain manually using a vibratome followed by clearing of the slice with RapiClear 1.52 (SunJin Lab) overnight at room temperature
4. mounting of the cleared sample with fresh RapiClear 1.52 reagent in a 0.25-mm-deep iSpacer microchamber (SunJin Lab).

We also imaged commercially available prepared slides containing rat testis tissues that were stained with hematoxylin and eosin (H&E, slide number 31-6464, Carolina Biological, Burlington, NC). This preparation is highly fluorescent and contains tissues with noticeable 3D structures.

### 2.2 Spinning disk setup

The system is based on an Olympus IX83 motorized inverted microscope. We used objective lenses UPLSAPO 20X/0.85 NA oil immersion, WD 0.17 mm, and UPLXAPO 60X/1.42NA oil immersion, WD 0.15 mm. Sample XY movements were controlled by a motorized stage (Applied Scientific Instrumentation, Eugene, OR, USA) while focusing was controlled by the IX83 microscope.

We used 445 nm (for GFP) and 532 nm (For H&E staining) lasers which were combined with dichroic mirrors and coupled into a multimode fiber (1 mm diameter, Thor Labs, Newton NJ, part number M35L02). To reduce speckle patterns from the multimode fiber, we shook the fiber with a home-made shaker based on a 120 mm computer fan. We imaged the end of the fiber on to the disk with a microscope objective, (Olympus 40X UPL) together with a 175 mm achromat lens (Edmund Optics, Barrington, NJ). This results in an evenly illuminated field, except for the remaining laser speckle [16]. The laser was reflected to the sample with a dual-band dichroic mirror (part ZT442/532rpc, Chroma, Bellows Falls, VT).

Fluorescence from the sample was imaged onto the spinning disk with the Olympus microscope and internal tube lens, then relayed onto a CMOS camera (Blackfly-S, Teledyne FLIR, Thousand Oaks, CA) with a pair of 175 mm focal length achromatic lenses (Edmund optics). A filter wheel (Lambda 10B, Sutter Instruments Novato, CA) housed the emission filters (ET500/50 for GFP and ET575/50 for H&E staining, Chroma).We used micro-manager software [7] to acquire the images with the 25MP camera. Micro-manager controls the camera, Olympus IX83 Z-axis drive, and filter wheel. Our setup is shown in Figure 1.

**Figure 1:**
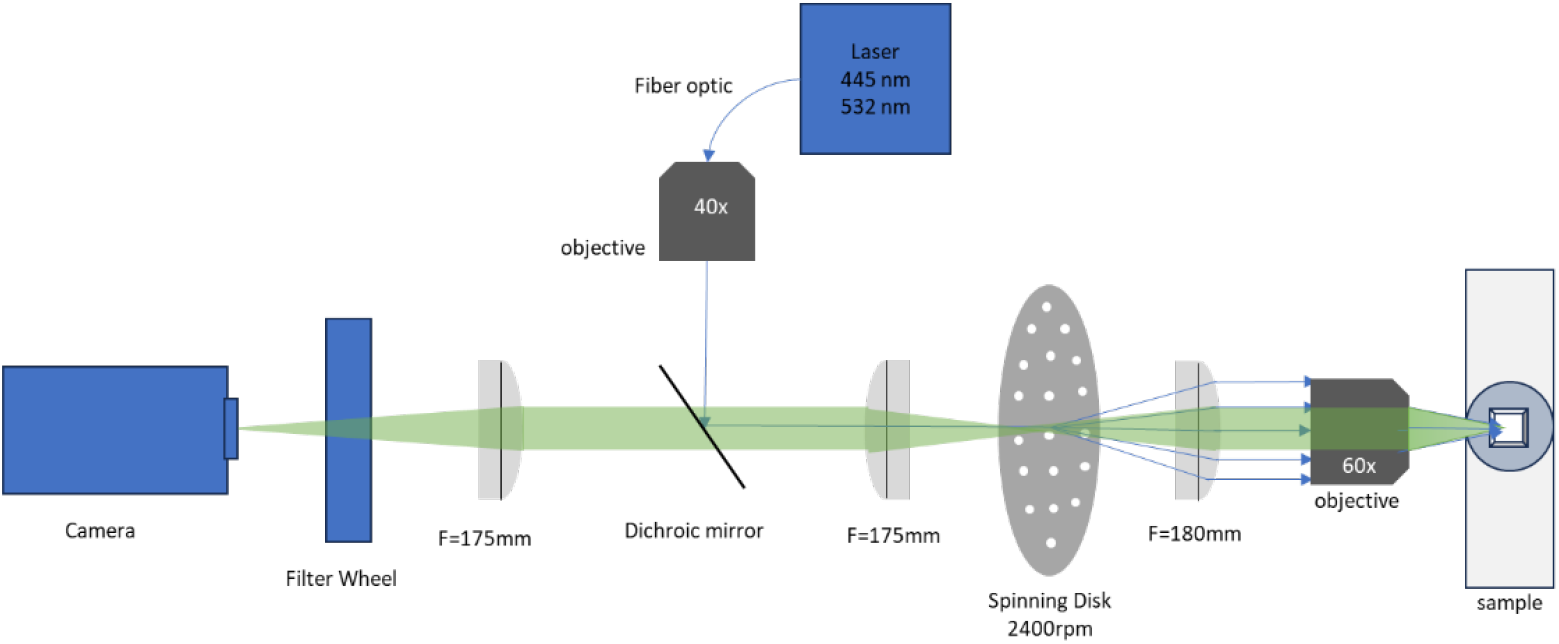
Setup for spinning disk confocal microscopy. A 40× objective and 175 mm lens image the end of a multimode fiber optic onto the spinning disk which is positioned in the primary image plane of the microscope.

Front range photomask (also known as Arizona Micro, Inc., Las Vegas, NV) produced the chrome-coated borosilicate glass disk based on a CAD file we provided. A Mathematica (Wolfram Research, Champaign, IL) program provided by Halpern, et al [14] was modified and used to produce the disk design. The photomask consists of a reflective chrome coating with an array of pinhole openings arranged in Archimedean spirals. The glass disk is mounted on a Stanford Research Systems (Sunnyvale, CA) model SR540 optical chopper. This was chosen because optical choppers are designed for constant rotation speed. The speed is constantly monitored and adjusted by the chopper using the timing notches. In our case, the timing notches were openings in the chrome photomask along the outer edge of the disk.

The disk design is based on the work of Halpern, et al. [14] with a few changes. Halpern et al. uses a computer hard disk drive as the motor, here we used the optical chopper. Halpern et al. has a disk design in which there are multiple sectors radially, so that different pinhole diameters and spacings can be accommodated on the same disk. We changed this to have only a single pinhole diameter and spacing on the disk. This means that we can use (nearly) 1:1 imaging throughout whereas Halpern et al. uses some extra magnification because each sector is smaller than the field of view of the camera chip. This means our optical setup is a bit larger (because of the focal lengths of lenses), but we have a nearly telecentric (no change in magnification when changing focus) design.

The optimal pinhole diameter in a spinning disk confocal microscope is given by [14] *d*= 1.22*λ* \**M/NA*, where λ is the wavelength of light, M is the magnification and NA is the numerical aperture. Table 1 shows the optimum pinhole size for a range of objective magnification and NA for a wavelength of 515 nm, e.g., for imaging GFP.

**Table 1:**
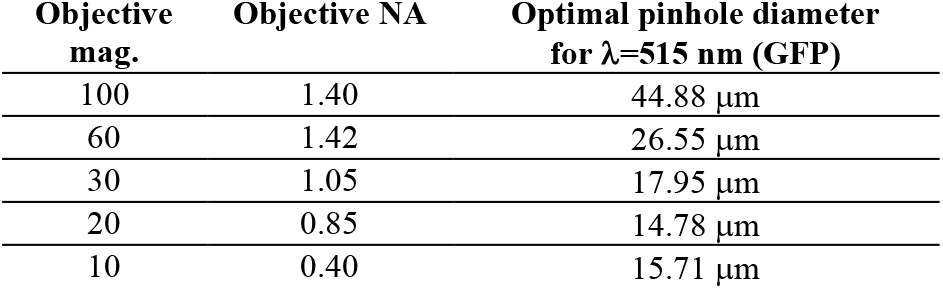
Optimal Pinhole size for Confocal Spinning Disk Microscopy at 515 nm.

Often, an inter-pinhole spacing of 10× the pinhole diameter is chosen [14]. In the final design, we chose a pinhole diameter of 18 μm and spacing of 180 μm. This was nearly optimal for a 30×/1.05 NA objective, but also a good compromise for use between 20× to 60× magnifications. Figure 2 shows a schematic of the disk design including the set of spirals (fig 2a), a single spiral (fig 2b), and an image from the camera when the disk is stopped (fig 2c). The sample in this case was a thin, uniform fluorescent film containing rhodamine dye.

**Figure 2:**
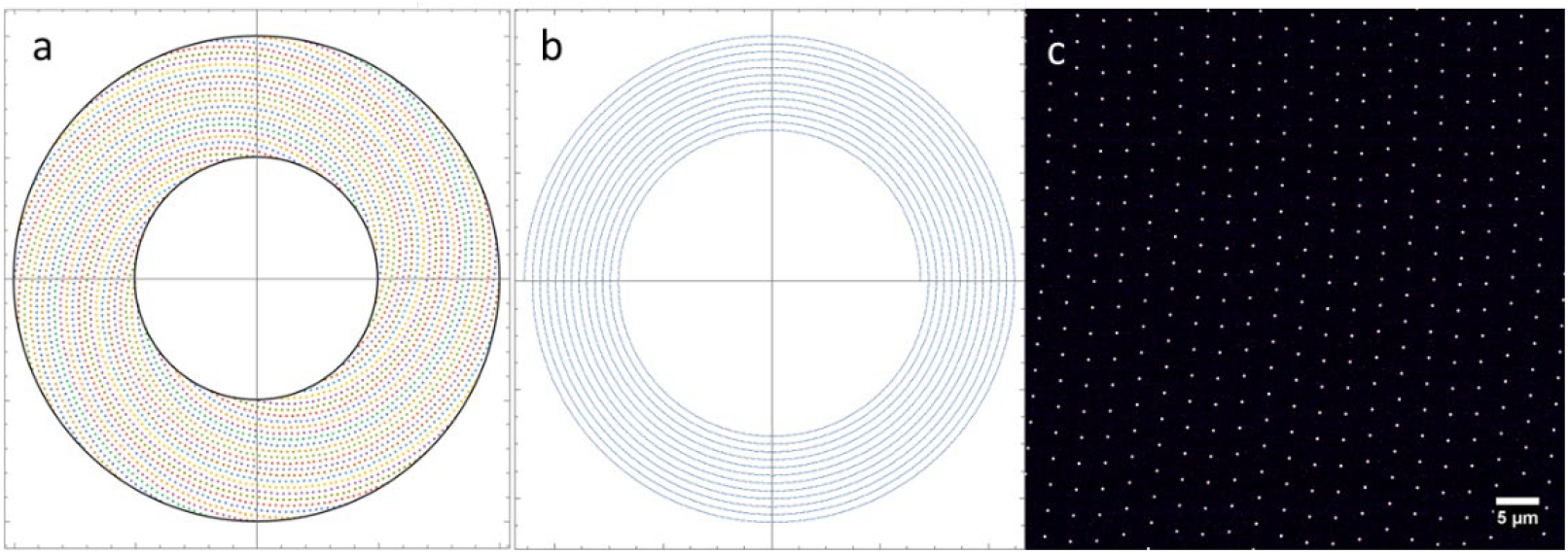
Schematic of spinning disk design. (a) set of spirals (b) single spiral (c) image from the camera when the disk is stopped showing pattern of pinholes in the middle of the sector as imaged by a 60x/1.42 oil immersion objective.

### 2.3 Machine vision camera

The sensor we used was the Sony IMX540 as implemented in the Blackfly S camera from Teledyne FLIR. This sensor is part of the Sony Pregius-S series of back-illuminated global shutter CMOS sensors. The IMX540, and its higher speed variant (IMX530) have been used in stereo vision applications [17] and for hyperspectral imaging [18]. An older Sony Pregius model (IMX250) was used for super-resolution single molecule localization microscopy (SMLM) [19]. With their low cost, CMOS cameras like the one used here are useful in multicamera setups such as the multi-plane structured illumination microscope used by our group [20]. Table 2 summarizes the parameters of the machine vision camera, and a camera more commonly used in spinning disk confocal microscopy, the Andor Zyla 4.2 scientific CMOS.

**Table 2.** Camera parameters and comparison.

| Camera | FLIR Blackfly S | Andor Zyla 4.2+ |
| --- | --- | --- |
| Part number | BFS-U3-244S8M-C | ZYLA-4.2P-CL10 |
| Sensor type | Sony IMX540 (Pregius S) | Andor Zyla |
| Quantum efficiency | 69% at 525 nm | 82% at 555 nm |
| Read noise | 2.31 e<sup>-</sup> | 0.9 e<sup>-</sup> |
| Dark current | 0.3 e<sup>-</sup>/pixel/sec | 0.1 e<sup>-</sup>/pixel/sec |
| Pixel size | 2.74 μm | 6.5 μm |
| Full well capacity | 9,648 e<sup>-</sup> | 30,000 e<sup>-</sup> |
| Pixel format | 5320 x 4600 (24.5 MP) | 2048 x 2048 (4.2 MP) |
| Maximum frame rate | 15 FPS | 100 FPS |
| Readout method | global shutter | rolling shutter |
| CMOS type | Back illuminated | Front illuminated |
| Sensor size | 14.599x12.626 mm<br>19.3 mm diag. | 13.3x13.3 mm<br>18.8 diag. |
| Interface | USB3 | Camera link |
| Weight of camera | 53 g | 1 kg |
| Size of camera | 29x29x39 mm | 133x80x82 mm |
| Approx. price | \$2,225 | \$18,000 |

This Sony IMX540 sensor has smaller pixels than are typically used in microscopy. This has the disadvantage that each pixel is less sensitive by the ratio of the pixel areas between the two cameras (6.5 μm^2^ vs. 2.74 μm^2^, a factor of 5.63). However, the advantage of a smaller pixel size is that the image will be oversampled at lower magnifications.

The resolution of a widefield fluorescence microscope is given by *d = 0.61 λ/NA*, where *d* is the distance between two objects that can just be resolved. To achieve proper sampling, we need at least two pixels covering the point spread function. That is, the expected resolution divided by the back-projected pixel size should be at least 2.0. Table 3 shows, for a variety of objectives, the expected resolution for λ = 515 nm, and the back-projected pixel size for the two different cameras (Sony IMX540 and Andor Zyla). The back-projected pixel size refers to the size of a camera pixel projected to the sample through a particular objective. For example, the Sony IMX540 sensor has a pixel size of 2.74 μm. When using at 10× objective, the back-projected pixel size would be 0.274 μm (274 nm). Table 3 also shows the sampling rate, calculated as the expected resolution divided by the back-projected pixel size. We can see that, in the widefield case, using the Sony IMX540 sensor allows oversampled imaging even at 10× magnification. Here we are using spinning disk confocal microscopy, which (when not using methods such as optical photon reassignment) does not typically achieve lateral resolution better than a widefield microscope [21].

**Table 3.** Sampling rate consideration for various objectives considering a widefield microscope:

| Objective<br>Mag/NA | Expected<br>Res. at 515<br>nm (nm) | Pixel size<br>FLIR (nm) | Sampling<br>rate (FLIR) | Over-<br>sampled?<br>(FLIR) | Pixel size<br>Andor (nm) | Sampling<br>rate<br>(Andor) | Over-<br>sampled?<br>(Andor) |
| --- | --- | --- | --- | --- | --- | --- | --- |
| 10X/0.40 | 785 | 274 | 2.87 | Yes | 650 | 1.21 | No |
| 20X/0.85 | 370 | 137 | 2.70 | Yes | 325 | 1.14 | No |
| 30X/1.05 | 299 | 91 | 3.28 | Yes | 216 | 1.38 | No |
| 40X/0.95 | 330 | 68.5 | 4.83 | Yes | 162.5 | 2.03 | Yes |
| 60X/1.35 | 233 | 45.6 | 5.10 | Yes | 108 | 2.14 | Yes |
| 60X/1.42 | 221 | 45.6 | 4.85 | Yes | 108 | 2.05 | Yes |
| 100X/1.40 | 224 | 27.4 | 8.19 | Yes | 65 | 3.45 | Yes |

### 2.4 Low-cost lasers

We used two lasers, a 2 W, 532 nm laser (Dragon laser, Changchun, China), and a 1.5 W, 447 nm laser (a low-cost laser acquired on the Chinese market through eBay). The lasers can be rapidly toggled on and off with 0-5 V signals, useful for toggling the lasers off in between camera exposures. We have found these lasers to be reliable and to have sufficient stability for our purpose. The higher powers are useful in this setup because we do not utilize the microlens-enhanced design as is found in commercial spinning disk units from Yokogawa [21]. The microlens-based design is a much more difficult construction because the microlenses must be positioned accurately above the pinholes and because a dichroic mirror must be positioned in between the two disks.

## 3. Results

### 3.1 Characterization of the camera

To characterize the two cameras, we imaged an intensity gradient pattern on an Argo-SIM slide (Argolight, Pessac, France). The intensity gradient pattern consists of 16 squares having different fluorescence intensity levels following a linear evolution. This is shown in Figure 3. The Andor Zyla camera has noticeably lower noise levels at 10 ms exposure. At 500 ms the images are more comparable.

**Figure 3:**
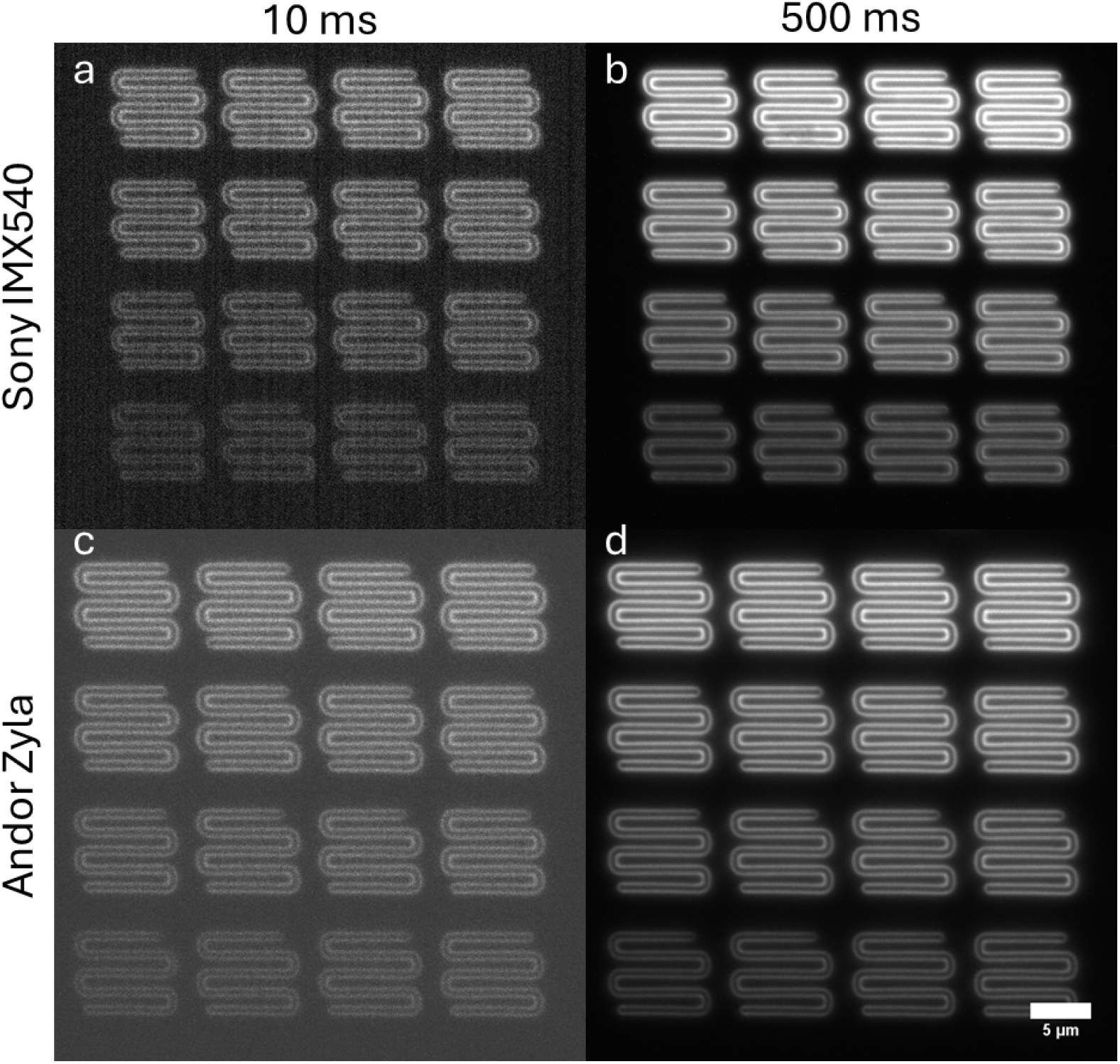
(a) 25 MP CMOS camera, 10 ms exposure, (b) 25 MP CMOS camera, 500 ms exposure. (c) Andor Zyla camera, 10 ms exposure, (d) Andor Zyla camera, 500 ms exposure. Objective: Olympus 100X/1.40 NA UPLSAPO oil immersion.

We then plotted the signal to noise ratio (SNR) for all 16 patches at 7 different exposure times ranging from 10 ms to 500 ms. We calculated SNR as the average intensity measured in each patch divided by the standard deviation of the background, obtained from a equal sized patch in the corner of the image in which there was no fluorescence signal. One can see that the Andor Zyla camera reached a higher SNR, with a maximum of 261.3 for the brightest patch at 500 ms exposure, compared with a SNR of 80.5 for this same patch imaged with the IMX540 CMOS camera at 500 ms exposure. At 500 ms exposure, the difference in SNR was consistent across the different brightness patches, with an average ratio of 3.48 +/-0.23, reflecting the difference in sensitivity between the two cameras (note this does not attempt to correct for the difference in pixel sizes between the cameras). This data is shown in figure 4.

**Figure 4:**
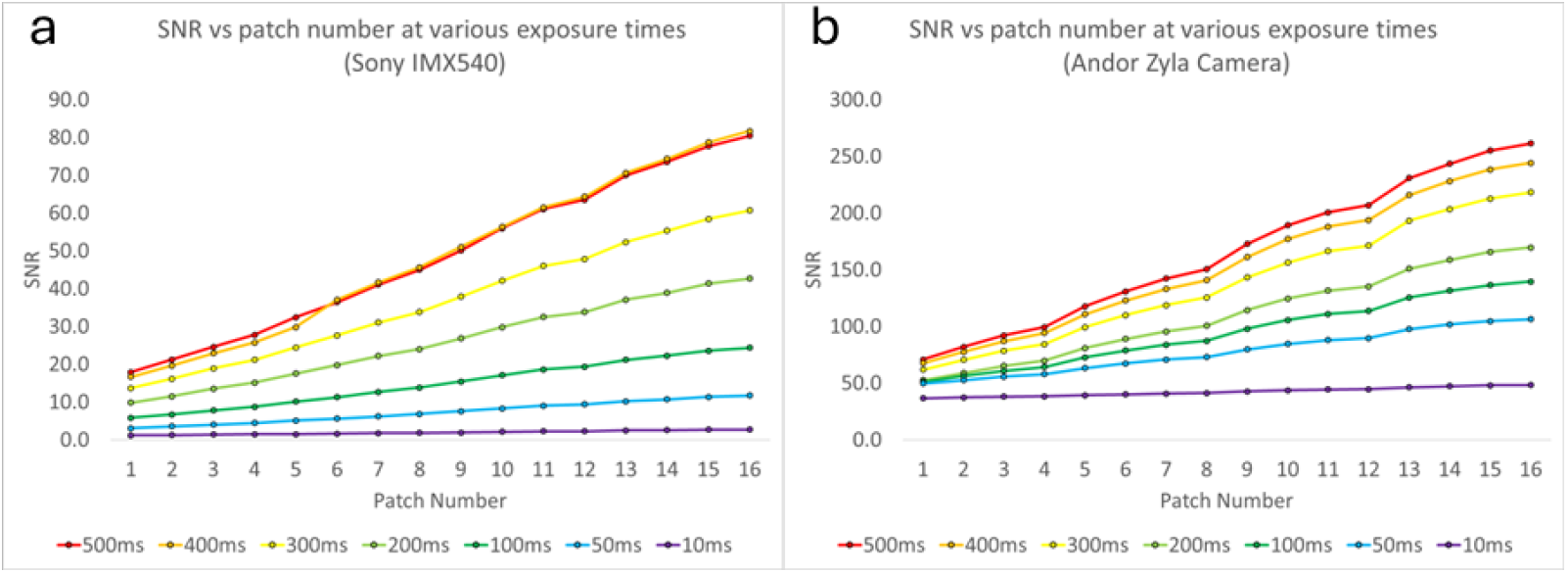
signal to noise ratio vs. patch number for various exposure times. (a) Sony IMX540. (b) Andor Zyla camera.

### 3.2 Imaging of biological tissues

We first imaged an optically cleared coronal mouse brain slice in which a subset of neurons express GFP [22]. This slice was matched to Paxinos and Franklin’s mouse brain atlas [23] to identify which section of the brain was being imaged. We matched or slice with slice 64 from the atlas. We further matched our sample to slice 92 of 132 in the Allen brain atlas [24,25]. Polynomial fits were made for both the horizontal and vertical directions using the edges and the central aqueduct as reference points. This allowed any point on this brain slice, recorded from the microscope stage coordinates, to be translated into the coordinates of the atlas for this particular slice. This method placed the neurons shown in figure 5 in the subiculum region of the hippocampus. Figure 5 shows a maximum intensity projection of 102 optical sections acquired with a spacing of 2 μm using an UplanSApo 20×/0.85 oil immersion objective (Olympus). This region is indicated in purple in figure 6, taken from the Allen brain atlas (Allen Mouse Brain Atlas, mouse.brain-map.org and atlas.brain-map.org).

**Fig 5:**
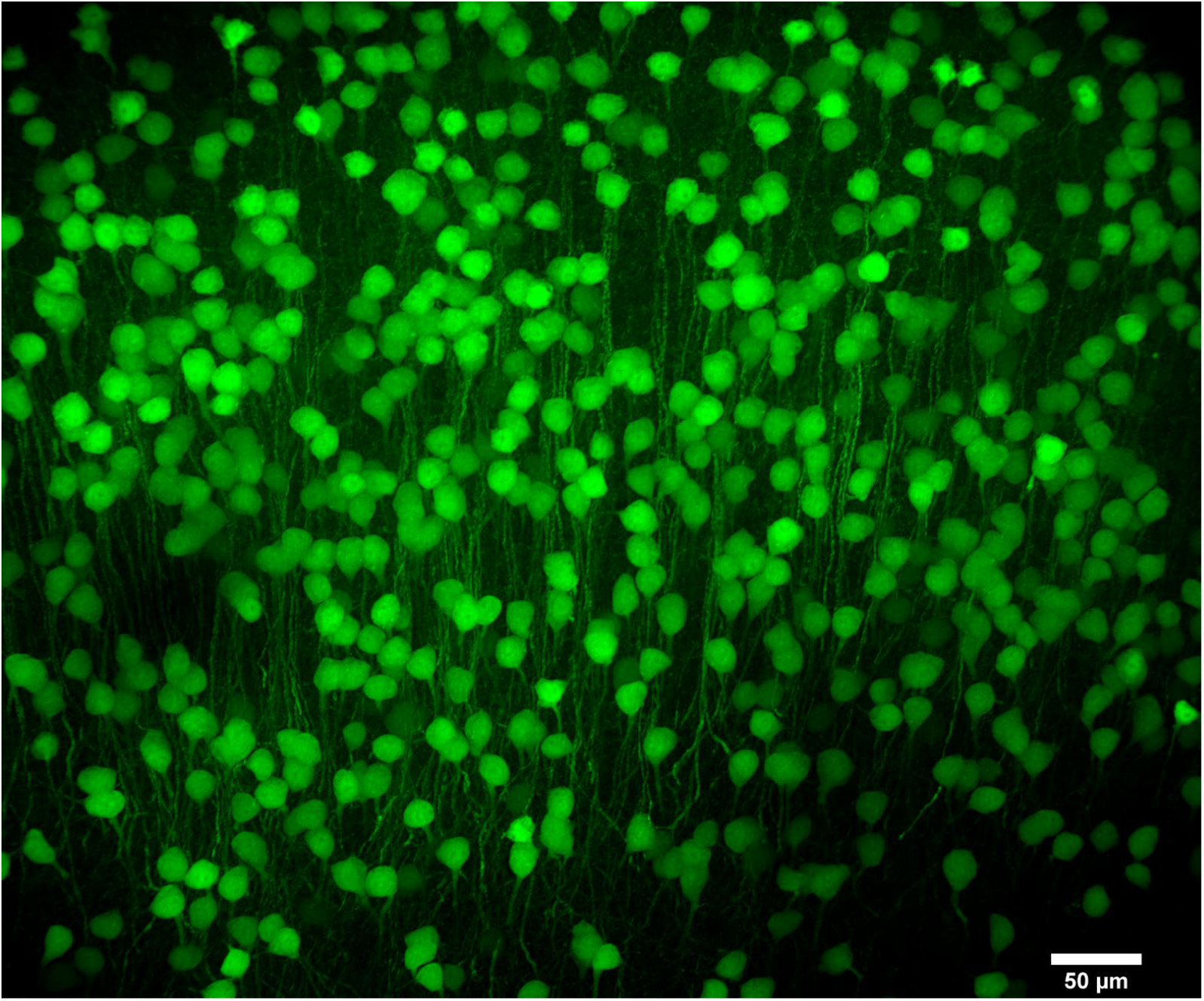
Neurons of the subiculum in the hippocampus of a *Thy1-GFP* mouse brain, spinning disk with 18 μm pinholes, Franklin and Paxinos slice 64. Allen brain atlas slice 92 of 132, 20X/0.85 NA oil immersion.

**Figure 6:**
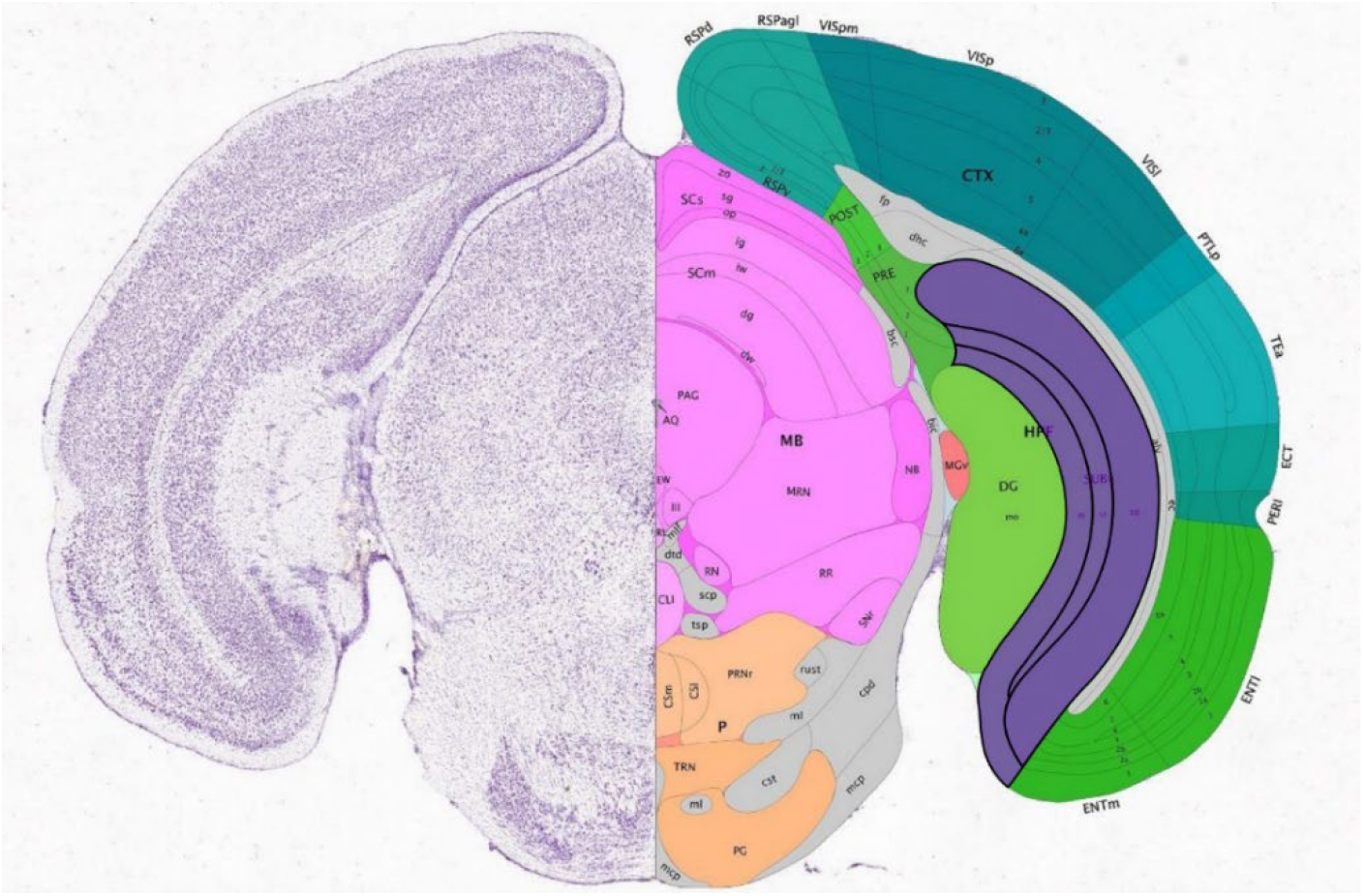
Allen brain slice 92 of 132 with subiculum highlighted in purple.

We also imaged commercially available prepared slides containing rat testis tissues that were stained with hematoxylin and eosin (slide number 31-6464, Carolina Biological, Burlington, NC). This preparation is highly fluorescent and contains tissues with intricate 3D structures. Figures 7 and 8 show two different areas of the sample and illustrate the high resolution imaging capabilities of the system over a large field of view.

**Fig 7:**
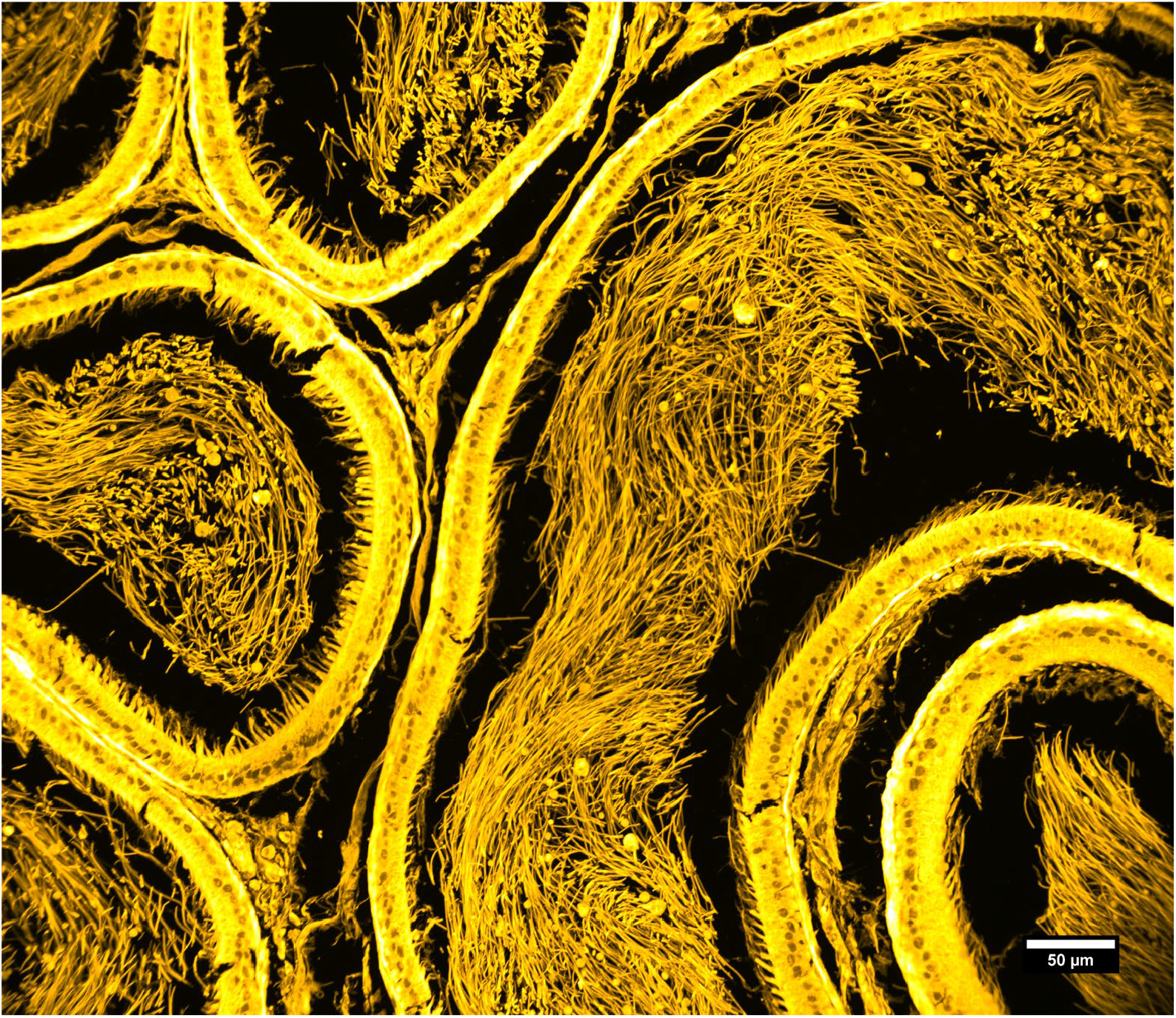
Rat testis, prepared slide. Maximum intensity projection of 69 slices, Z-spacing 0.5 um. Exposure time 100 ms. Objective: 20×/0.85 NA oil immersion.

**Figure 8:**
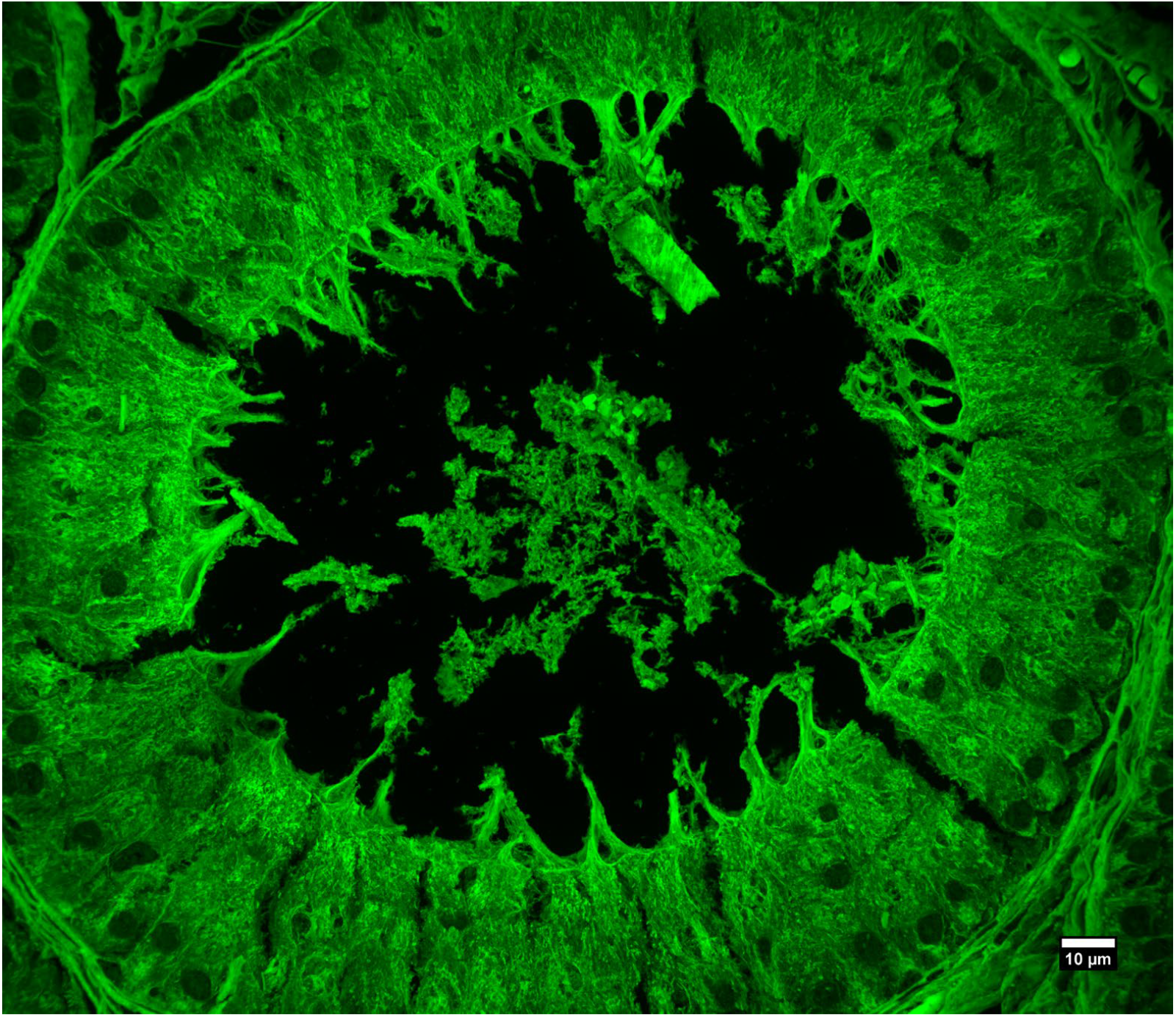
Rat testis, prepared slide. Maximum intensity projection of 72 slices, Z-spacing 150 nm, Exposure time 200 ms. Objective: 60×/1.42NA oil immersion.

## 4. Discussion

Lower cost designs for advanced fluorescence microscopy applications such as spinning disk confocal microscopy are expected to increase their accessibility and use by a greater number of researchers. In particular, the higher cost of scientific CMOS cameras has limited their use in multi-camera setups, and has sometimes led to complex optical arrangements in order to direct the multiple images onto one camera. Some examples where multiple images were routed onto a single camera include certain implementations of the programmable array microscope [26,27] and the dual objective microscope for single molecule localization microscopy [16,28]. Even very recent designs have faced this issue [29]. Typically, the one camera approach causes the need for a reimaging system, which inevitably increases optical aberrations, particularly axial chromatic aberration.

The 25 megapixel camera used here together with the spinning disk offers large, highly detailed, optically sectioned images that are particularly appealing when viewed on modern high resolution computer monitors. Image stitching methods would usually be required to achieve the same result.

## Author Contributions

T.C.P.: acquired the data, analyzed the data, and helped design the spinning disk; S.L; J.R.H.: acquired the data and analyzed the data; B.L., analyzed data; G.M.H.: conceived the project, acquired the data, analyzed the data, supervised the research, and wrote the paper. All authors have read and agreed to the published version of the manuscript.

## Funding

The research reported in this publication was supported by the National Institute of General Medical Sciences of the National Institutes of Health under award number 2R15GM128166-02. This work was also supported by the UCCS BioFrontiers center.

## Notes

### Competing Interest Statement

The authors have declared no competing interest.

